# Cell size overrides ganglion mother cell fate specification in Drosophila

**DOI:** 10.64898/2026.09.08.750127

**Authors:** Inès Jmel Boyer, Nicolas Loyer, Madeleine Davies, Isla Mortimer, Chantal Roubinet, Hedda A. Meijer, Kim Dale, Jens Januschke

## Abstract

The size of cells varies enormously in biology and how cell size and function are linked are fundamental questions. Asymmetrically dividing *Drosophila* neural stem cells, also called neuroblasts, produce a large neuroblast and a smaller ganglion mother cell (GMC) at each division. However, why GMCs need to be smaller than neuroblasts has largely remained unexplained. Here, we developed acute and reversible methods to alter daughter-cell size during a single neuroblast division while preserving cortical polarity and asymmetric inheritance of fate determinants. We identify atypical protein kinase C (aPKC) as a regulator of size asymmetry and show that transient and partial co-inhibition of aPKC and Rho kinase (Drok in *Drosophila)* generates daughters spanning a size range, including size-wise near-symmetric sibling cells while polarity is preserved. We find that above a certain size threshold daughter cells retain neuroblast-like properties despite inheriting GMC determinants: they divide repeatedly, exhibit neuroblast-like cell-cycle timing, and undergo size-wise asymmetric division. Smaller daughters below this threshold instead follow a GMC-like trajectory. Thus, in Drosophila type I neuroblasts, daughter-cell size can act as a regulator of lineage behaviour, overriding inherited fate cues in sufficiently large daughter cells.

## INTRODUCTION

How cells regulate their size and how cell size relates to cell function and fate are fundamental questions in developmental biology (Amodeo and Skotheim 2016). Cell size is influenced by growth-regulatory pathways including PI3K–AKT–mTOR, Myc and Hippo, as well as by mechanisms coupling growth to cell-cycle progression and division (Lloyd 2013). However, another way cells can regulate size is by altering the position of the cytokinetic furrow during cell division which results in size-wise different daughter cells. This type of division plays a role in development where the differences in daughter cell size are often correlated with differences in fate or function (Morin and Bellaïche 2011; Delgado and Cabernard 2020).

There are clear cases in which cell size is instructive for fate following asymmetric division. In the green algae Volvox unequal embryonic divisions produce large gonidial initials and small somatic initials and cell size resulting from eccentric cleavage is an important developmental input for fate (Kirk et al. 1991; Kirk et al. 1993). During development, cell size changes can also act as a cell fate switch. In the stomatal lineage of Arabidopsis meristemoids undergo repeated asymmetric self-renewing divisions through which they get smaller. A fate switch occurs and stem cells commit to terminal differentiation when stem cells cross a critical cell-size threshold (Gong et al. 2023). Cell size thresholds also matter during embryogenesis of *C.elegans* where cell size limits polarity and therefore asymmetric division potential in the P-lineage. Below a size threshold, polarity becomes unstable as reducing P-lineage cell size genetically or physically causes loss of polarity and a premature switch from asymmetric to symmetric divisions (Hubatsch et al. 2019).

Anisotropic Myosin activity has been found to drive eccentric positions of the cytokinetic furrow in the Q neuroblast lineage of *C.elegans* and in *Drosophila* neuroblasts (Cabernard et al. 2010; Ou et al. 2010). *Drosophila* neuroblasts divide asymmetrically and at each mitosis they polarize the PAR complex including Par3/Bazooka (Baz) and atypical protein kinase C (aPKC) at the apical pole which leads to the localization of fate determining molecules to the basal pole. Fate determining molecules include the transcription factor Prospero (Pros) and the translational repressor Brain Tumour (Brat) which both rely on Miranda (Mira) for asymmetric segregation, as well as the Notch signalling regulator Numb. The spindle then aligns with this polarity axis resulting in unequal segregation of molecules: the PAR complex is inherited by the larger self-renewed neuroblast, and the basally localized factors end up in the smaller daughter cell, called ganglion mother cell (GMC) (Gallaud et al. 2017; Delgado and Cabernard 2020; Loyer and Januschke 2020).

In the *Drosophila* larval central brain, neural stem cells are broadly classified as type I and type II neuroblasts, which differ mainly in the proliferative behaviour of their daughter cells (Boone and Doe 2008; Bayraktar et al. 2010). Type I neuroblasts divide asymmetrically to self-renew while producing a GMC, which typically undergoes a single terminal division to generate two neurons or glial cells. In contrast, type II neuroblasts generate intermediate neural progenitors (INPs) which mature and undergo several rounds of asymmetric division to produce multiple GMCs (Bello et al. 2008; Bowman et al. 2008).

Both type I and type II neuroblast lineages use asymmetrically segregated fate determinants, including Brat and Numb, to restrict self-renewal in their daughter cells. In type I lineages, Pros acts as a binary switch in the GMC, repressing stem-cell genes and activating differentiation genes, and GMCs undergo one terminal division (Choksi et al. 2006). Brat, a sequence-specific RNA-binding post-transcriptional (Laver et al. 2015), regulates neuroblast identity factors including *deadpan* (Reichardt et al. 2018), and its RNA-binding ability is essential for its tumour-suppressor function (Connacher et al. 2026). In type II lineages, whose neuroblasts lack Pros, Brat and Numb promote immature intermediate neural progenitor (INP) maturation and prevent reversion to neuroblast identity (Bowman et al. 2008). Mature INPs then undergo multiple asymmetric divisions to generate GMCs. Numb acts cell-autonomously to inhibit Notch signalling in the daughter cell that inherits it (Frise et al. 1996). Therefore, asymmetric fate-determinant segregation is the best-established mechanism to explain how neuroblasts and differentiating daughter cells are established during asymmetric division.

Neuroblasts and their GMC daughters differ also considerably in size. Although the mechanisms that generate this size asymmetry are relatively well understood (Connell et al. 2011; Roth et al. 2015; Roubinet et al. 2017; Tsankova et al. 2017; Pham et al. 2019; Montembault et al. 2023; Loyer et al. 2026b), the functional significance of producing a GMC that is much smaller than its neuroblast sibling remains unclear. Intriguingly, GMC size is relatively constant in embryonic and larval lineages alike (Fuse et al. 2003; Homem et al. 2013; Loyer et al. 2026b), suggesting that GMC size may be functionally constrained. By contrast, neuroblast size can decrease over developmental time, with corresponding effects on cell-cycle length (Fuse et al. 2003; Homem et al. 2013). This raises the possibility that GMCs must remain below a maximum size to undergo their characteristic terminal size-wise symmetric division, whereas neuroblasts must remain above a minimum size to retain their proliferative capacity.

Mutant analyses support a link between physical asymmetry and lineage outcome in *Drosophila* neuroblasts, but the specific contribution of cell size remains unclear. In *dlg gβ13F* double mutants, neuroblasts lose normal polarity and size asymmetry, with context-dependent consequences: embryonic neuroblasts initially divide but later differentiate into neurons, producing clones of homogeneous temporal identity, whereas larval neuroblasts overproliferate (Kitajima et al. 2010). Because both polarity and size asymmetry are disrupted, these phenotypes cannot be attributed specifically to daughter-cell size. *gβ13F* single mutants largely retain cortical polarity and fate-determinant segregation but produce nearly equal-sized daughters; repeated near-symmetric divisions cause lineage alterations including neural defects, suggesting that eccentric division helps maintain neuroblast stem-cell properties (Fuse et al. 2003). However, Gβ13F also regulates spindle geometry and G-protein signalling, so cell size is not altered in isolation. Manipulating myosin-dependent contractility similarly changes daughter-cell size asymmetry (Tsankova et al. 2017) and can increase neuroblast number and disrupt lineage outcomes (Delgado et al. 2025). Yet both genetic and contractility-based manipulations are chronic and potentially pleiotropic. Thus, it remains unresolved whether daughter-cell size itself influences neuroblast self-renewal or GMC differentiation when fate-determinant segregation is intact, or whether a size threshold governs fate decisions.

Here, we developed experimental approaches that allow acute manipulation of daughter-cell size during a single neuroblast division while preserving asymmetric fate-determinant segregation. We identify aPKC as a regulator of GMC size and show that mild, transient co-inhibition of aPKC and Rho kinase (Drok) generates daughter cells spanning a broad range of sizes, including near-symmetric siblings, while polarity and determinant segregation remain asymmetric. Following inhibitor washout, sufficiently large determinant-inheriting daughters undergo repeated asymmetric divisions with neuroblast-like cell-cycle timing, whereas smaller daughters display GMC-like behaviour. These findings reveal daughter-cell size as an important determinant of subsequent lineage behaviour and suggest that eccentric cytokinesis normally positions sibling cells on opposite sides of a relevant size threshold.

## RESULTS

### Novel tools for specific and acute inhibition of Rho kinase in *Drosophila*

Cell size asymmetry in neuroblasts depends on the spatiotemporal regulation of non-muscle Myosin II (Myosin) activity (Connell et al. 2011; Roubinet et al. 2017; Montembault et al. 2023; Loyer et al. 2026a). To manipulate Myosin activity acutely and reversibly, we sought alternatives to Blebbistatin, which is ineffective in *Drosophila* cells (Straight et al. 2003). Because Myosin activity depends on Rho kinase and the commonly used Rho kinase inhibitor Y-27632 has limited specificity (Atwood and Prehoda 2009), we developed two approaches to inhibit *Drosophila* Rho kinase (Drok). First, we generated the analog-sensitive *drok^as2^* allele and mutated the gatekeeper residue (Bishop et al. 2000) M164 to Alanine by CrispR yielding viable and fertile flies. This method allows acute and specific and reversible inhibition of kinases (Lopez et al. 2014). Second, we tested BAY549 (also known as TC-S-7001), a Rho kinase inhibitor developed for vertebrate kinases (Schirok et al. 2008), and found that it also inhibits Drok. *In vitro*, both approaches potently inhibited Drok activity (using a recombinant constitutively active version, DrokCAT, *see methods*): the IC_50_ values for DrokCAT^as2^ with 1-NAPP1 and DrokCAT^WT^ with BAY549 were <3 nM, compared with ∼0.8 µM for DrokCAT^WT^ with Y-27632 (**Figure 1A**, **Figure 1 supplement 1**). Thus, in this assay, BAY549 and the analog-sensitive DrokCAT^as2^/1-NAPP1 combination inhibited Drok at least two orders of magnitude more potently than Y-27632.

**Figure 1.**
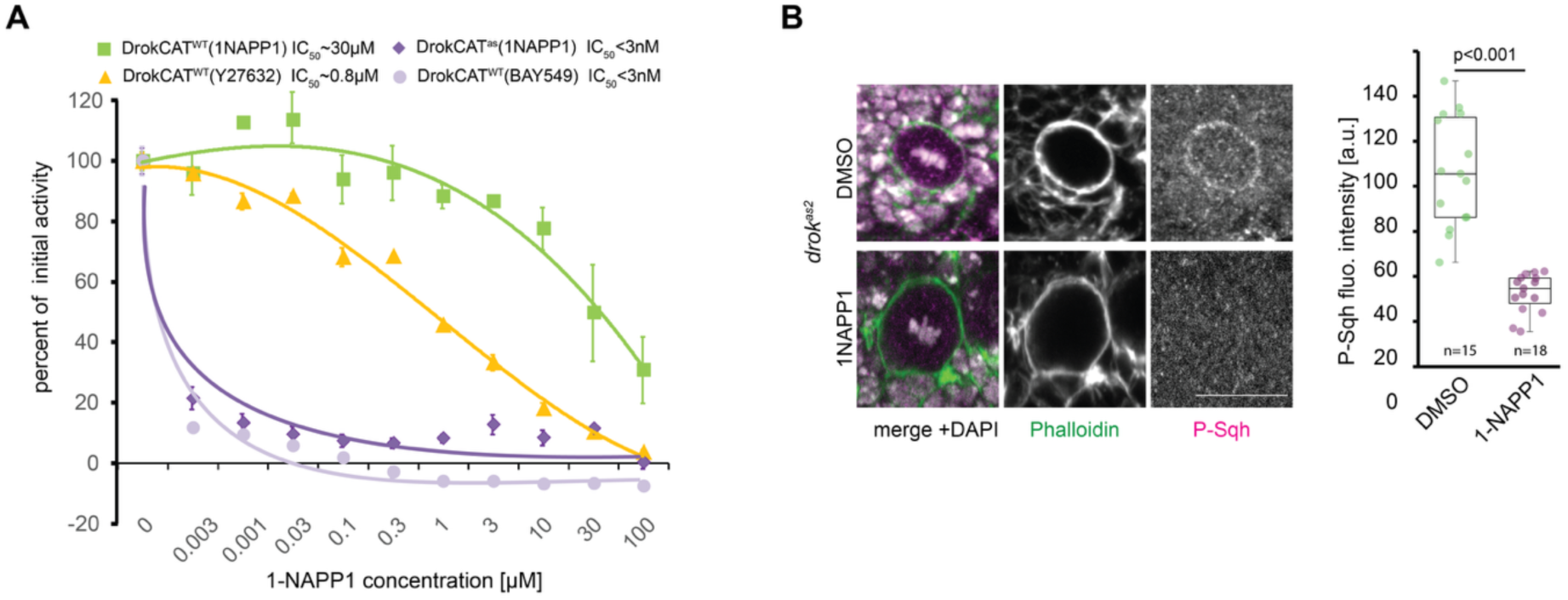
Acute and specific inhibition of Drok by BAY54S and drok^as2^ / 1-NAPP1. **A**) *In vitro* kinase assay to determine the IC50 for DrokCAT^WT^ with 1-NAPP1, Y-27632 or BAY549 and DrokCAT^as2^ with 1-NAPP1 (*see methods*). **B**) Fixed whole mount brain stained for the indicated factors measuring the ability of acute Drok inhibition using the *drok^as2^* allele and 20µM 1-NAPP1 to reduce phosphorylation of MRLC, scale bar 10µm, quantification to the right.

We also performed experiments to establish that both *drok^as2^*/1-NAPP1 and BAY549 disrupt Myosin-dependent functions *in vivo* (**Figure 1 supplement 2**). For instance, examining the phosphorylation of the myosin regulatory light chain (MRLC), a conserved Rho kinase substrate (Amano et al. 1996; Winter et al. 2001). MRLC Ser20 is phosphorylated in mitotic larval neuroblasts (Tsankova et al. 2017), and Drok inhibition using *drok^as2^*/1-NAPP1 significantly reduced this signal (**Figure 1B**).

Together, these results show that BAY549 and *drok^as2^*/1-NAPP1 provide effective approaches for acute Drok inhibition both *in vitro* and *in vivo* and indicate that Drok is a major kinase responsible for MRLC Ser20 phosphorylation in mitotic neuroblasts.

### Acute Drok inhibition disrupts cytokinesis and nuclear partitioning and affects Sqh localisation

We then examined the effects of Drok inhibition on neuroblast division (**Figure 2**). We first performed a dose-response experiment to increasing levels 1-NAPP1 to inhibit of Drok using bright-field microscopy and *drok^as2^* neuroblasts in primary culture. Increasing concentrations of 1-NAPP1 progressively reduced the frequency of what appeared to be size-wise asymmetric divisions and increased the frequency of cytokinesis failure. At intermediate concentrations, we also observed an increase in an unusual division outcome in which both nuclei were inherited by the larger neuroblast while a smaller, daughter cell consequently without a nucleus (termed ‘anucleated) was generated. These small anucleated cell-like structures were stable and able to generate a cortical microtubule network (**MOV2**). Inhibiting Drok caused also what appeared enlarged GMCs, binucleated neuroblasts and we observed lagging chromosomes which preceded cytokinetic failure (**Figure 2 supplement 1A,B,C**). We further analysed the effect of Drok inhibition on neuroblast division in whole mount brains using BAY549 and observed cytokinesis failure, binucleated neuroblasts with anucleated daughter cells at similar frequencies as in culture (**Figure 2 supplement 2A**).

**Figure 2.**
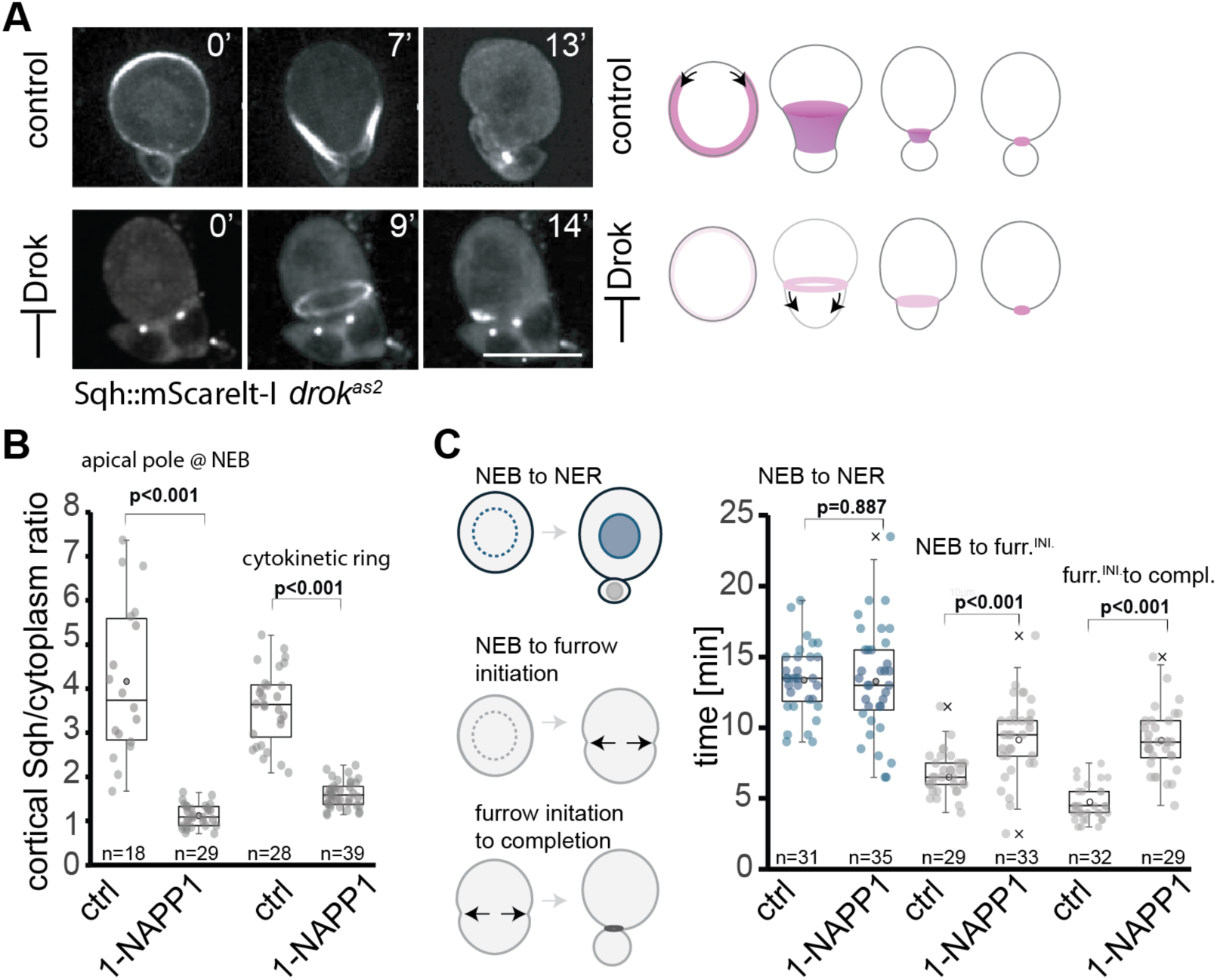
Sqh localisation changes and effects on the timing of neuroblast division upon strong Drok inhibition. **A**) Selected images from a time series of *drok^as2^* larval neuroblasts in primary cell culture expressing Sqh::mScarlet-I to label myosin upon addition of DMSO (control, top row) or 20µM 1-NAPP1 (bottom row), schematic of the observed effects shown on the right, scale bar 10µm. **B**) Quantification of cortical Sqh::mScarlet-I levels at the apical pole and in the cytokinetic ring comparing control and Drok inhibition via *drok^as2^*/ 20µM 1-NAPP1. **C**) Quantification of the length of mitosis and the duration of cell division. Schematics indicating what was measured to the left (nuclear envelope breakdown, NEB; nuclear envelope reformation, NER), Statistical tests: Mann-Whitney U test.

The reduction in MRLC Ser20 phosphorylation following Drok inhibition (**Figure 1B**) suggested that Myosin recruitment to the neuroblast cortex might also be affected. Consistent with this, strong Drok inhibition markedly reduced cortical Sqh levels during mitosis (**Figure 2A,B**). Sqh nevertheless accumulated in the cytokinetic ring, although at significantly lower levels than in controls, and the ring remained capable of contraction. Drok inhibition also altered the timing of cytokinesis. Compared with controls, furrow ingression began later and took longer to complete, although constriction consistently proceeded towards the basal side of the cell (**Figure 2A,C**; **MOV3**). In contrast, the duration of mitosis, measured from nuclear envelope breakdown to nuclear envelope reformation, was unchanged (**Figure 2C**), which we also observed in BAY549 treated neuroblasts in whole mount brains (**Figure 2 supplement 2B**).

Together, these observations show that strong acute Drok inhibition compromises both cytokinesis and nuclear partitioning during larval neuroblast divisions. They also show that Drok activity is required for efficient cortical recruitment of Myosin and for normal accumulation of Myosin at the cytokinetic furrow. Reduced Myosin levels do not prevent furrow contraction but delay its onset and prolong cytokinesis.

### Neuroblast polarity is maintained despite strongly reduced cortical Myosin

We next examined the temporal dynamics of apical Sqh recruitment during neuroblast division. In control cells, oscillatory apical Sqh recruitment was detectable approximately 20 min before nuclear envelope breakdown (NEB), followed by a further increase in cortical Sqh immediately before NEB and subsequent apical enrichment (Roubinet et al. 2017). All these features were strongly reduced or absent following Drok inhibition (**Figure 3A,A′**), confirming that Drok activity is required for efficient recruitment of Myosin to the neuroblast cortex.

**Figure 3.**
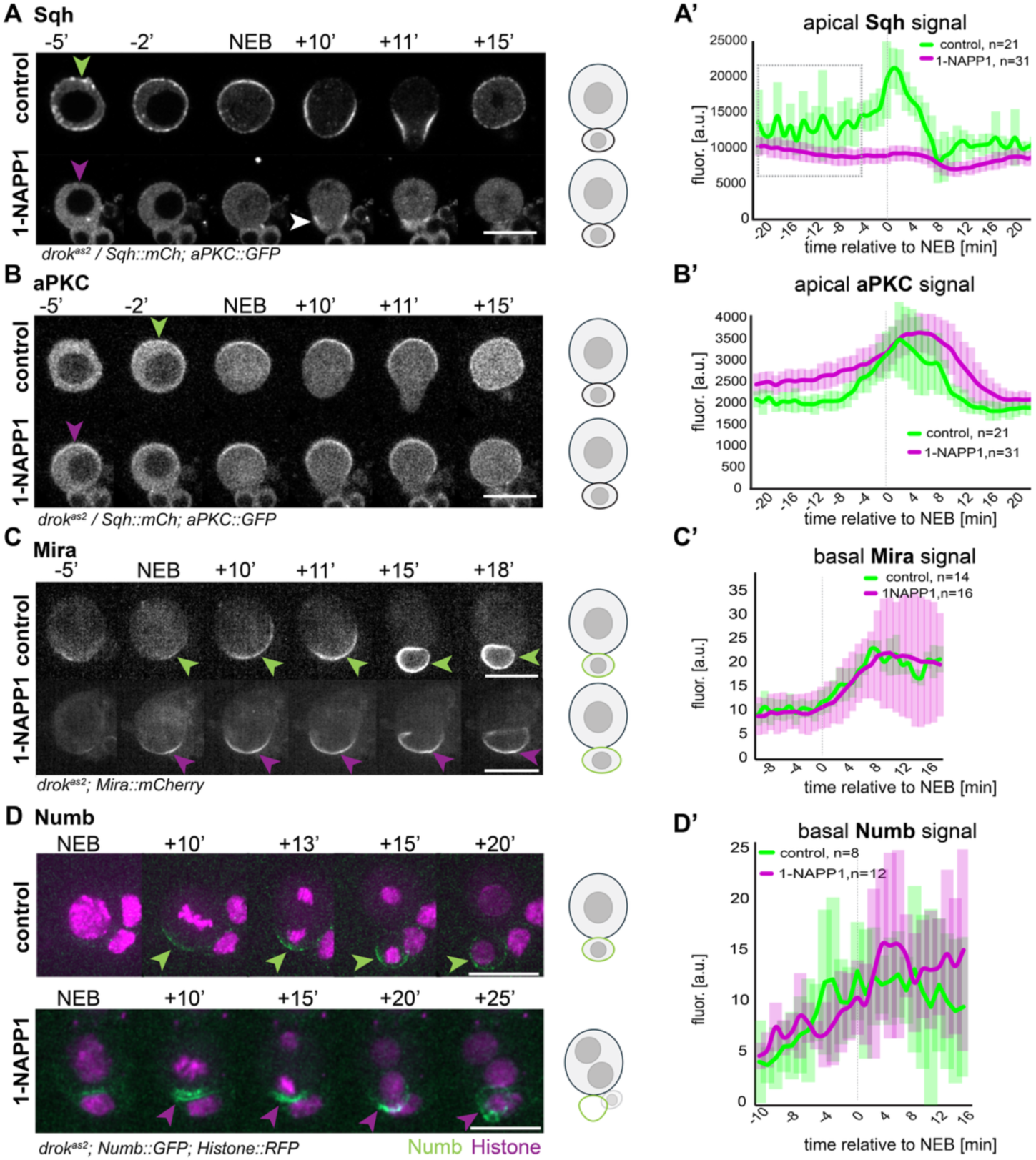
Neuroblast polarity and cell fate determinant localisation upon strong Drok inhibition. **A**) Time series of larval *drok^as2^* neuroblasts in primary cell culture expressing Sqh::mScarlet-I of control (DMSO treated) or upon 20µM 1-NAPP1 treatment. **A’**) Schematic of division outcome and quantification of apical Sqh to the right. **B**) The same cell as in A but showing the aPKC::GFP. **B’**) Schematic of division outcome and quantification of apical aPKC to the right. **C**) Time series of larval *drok^as2^* neuroblasts in primary cell culture expressing Mira::mCherry of control (DMSO treated) or upon 20µM 1-NAPP1 treatment. **C’**) Schematic of division outcome and quantification of basal Mira to the right. **D**) Time series of larval *drok^as2^* neuroblasts in primary cell culture expressing Numb::GFP of control (DMSO treated) or upon 20µM 1-NAPP1 treatment. **D’**) Schematic of division outcome and quantification of basal Numb to the right (note that Numb::GFP is expressed endogenously at low levels yielding noisy data).

Because Myosin has been implicated in neuroblast polarisation (Barros et al. 2003; Hannaford et al. 2018), we next asked whether this strong reduction in cortical Sqh affected polarity. Despite reduced cortical Myosin, neuroblast polarity was largely maintained. aPKC whose localisation depends on Baz (Wodarz et al. 2000), as well as the basal proteins Mira and Numb, all acquired asymmetric cortical localisation following Drok inhibition (**Figure 3**). Drok inhibition nevertheless altered the dynamics of some polarity components. aPKC appeared at the cortex earlier than in controls and persisted for longer during division (**Figure 3B,B′)**. Mira showed broadly similar recruitment kinetics and peak levels, although basal Mira levels became more variable approximately 10 min after NEB, most likely a consequence of the variable outcomes of cytokinesis under these conditions (**Figure 3C,C′**). Numb also remained asymmetrically localised and segregated into the smaller daughter compartment, even when the corresponding nucleus was retained by the larger daughter cell (**Figure 3D,D′**; **MOV4**).

Thus, strong Drok inhibition substantially reduces cortical Myosin recruitment without abolishing neuroblast polarity or asymmetric fate-determinant segregation consistent with recent findings (Gujar et al. 2025). The prolonged cortical persistence of aPKC further coincides with the delayed cytokinetic-ring contraction observed under these conditions.

### aPKC regulates GMC size and partial co-inhibition of aPKC and Drok strongly increases GMC size variability

We previously found that treatment with 25 µM Y-27632 produces enlarged GMCs that continue to inherit Mira (Hannaford et al. 2018). However, at concentrations commonly used in *Drosophila*, Y-27632 can inhibit both Drok and aPKC (Atwood and Prehoda 2009). We therefore examined the contribution of each kinase to daughter-cell size asymmetry separately, using absolute GMC size rather than NB/GMC size ratio as a quantitative readout (Loyer et al. 2026b). Strong inhibition of aPKC by 10µM 1-NAPP1 significantly increased GMC size (**Figure 4 A,B**). However, while partial inhibition of Drok with 1µM 1-NAPP1 or aPKC resulted only in a nonsignificant increase in GMC size, partial co-inhibition of both kinases at that concentration did significantly increase it (**Figure 4C,D**, **MOV5**).

**Figure 4.**
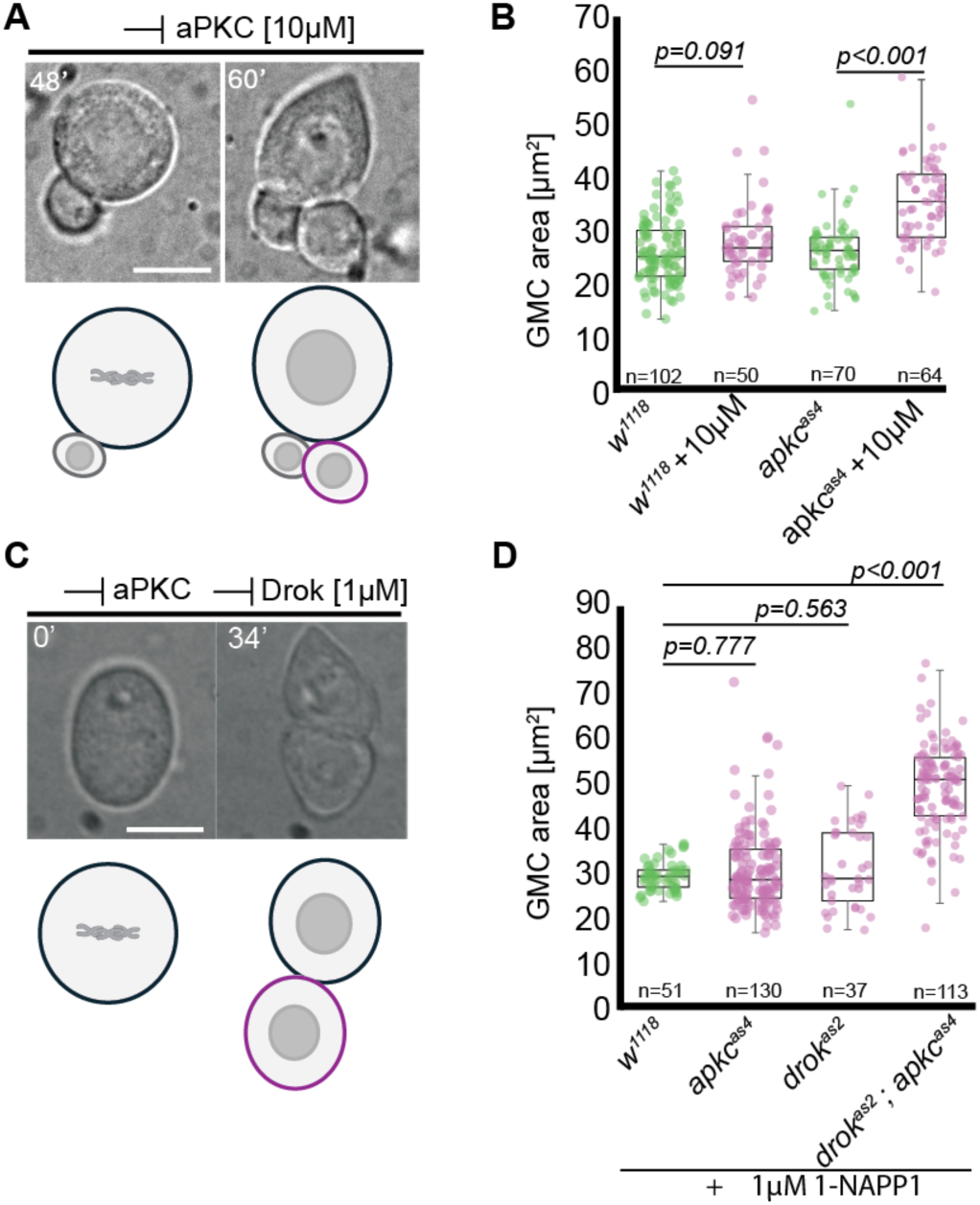
Partial aPKC and Drok co-inhibition significantly impacts GMC size. **A**) Panels from a DIC bright field time series of an *apkc^as4^* neuroblast in primary cell culture upon the addition of 10µM 1-NAPP1. Schematic of phenotype below. **B**) Quantification of GMC cell size using the ‘equatorial’ area of the GMC directly after cytokinesis under the indicated conditions. **C**) Panels from a DIC bright field time series of a *drok^as2^ apkc^as4^* double mutant neuroblast in primary cell culture upon the addition of 1µM 1-NAPP1. Schematic of phenotype below. **D**) Quantification of GMC cell size upon addition of 1µM 1-NAPP1 for the indicated genotypes. Scale bars: 10µm. Statistical tests: Mann-Whitney U test.

Across four independent experiments observing 164 neuroblasts upon partial co-inhibition of aPKC and Drok, we observed ∼27% of cytokinetic failures, ∼12% of binucleated neuroblasts with an anucleated daughter cell, ∼40% divisions with a clear size asymmetry, although about one third of those appeared to have enlarged GMCs (see also Figure 6) and ∼21% appeared to undergo near symmetric divisions (**MOV6**).

These results identify aPKC as a regulator of daughter-cell size asymmetry and show that partial co-inhibition of aPKC and Drok has a stronger effect on GMC size and the fidelity of cytokinesis than inhibition of either kinase alone.

### Polarity is maintained upon partial aPKC and Drok co-inhibition

We next asked whether the strong reduction in daughter-cell size asymmetry caused by partial co-inhibition of Drok and aPKC was accompanied by defects in neuroblast polarity or fate-determinant segregation. We first analysed whole mount brains treated with Colcemid to enrich for neuroblasts in metaphase. We also treated brains for 45min with 1µM 1-NAPP1 before fixation to capture more stages of division. Across these conditions, aPKC, Mira, Pros and Numb retained asymmetric localisation following partial co-inhibition (**Figure 5A–D**). In Colcemid arrested neuroblasts Mira showed stronger basal intensities than in controls.

**Figure 5.**
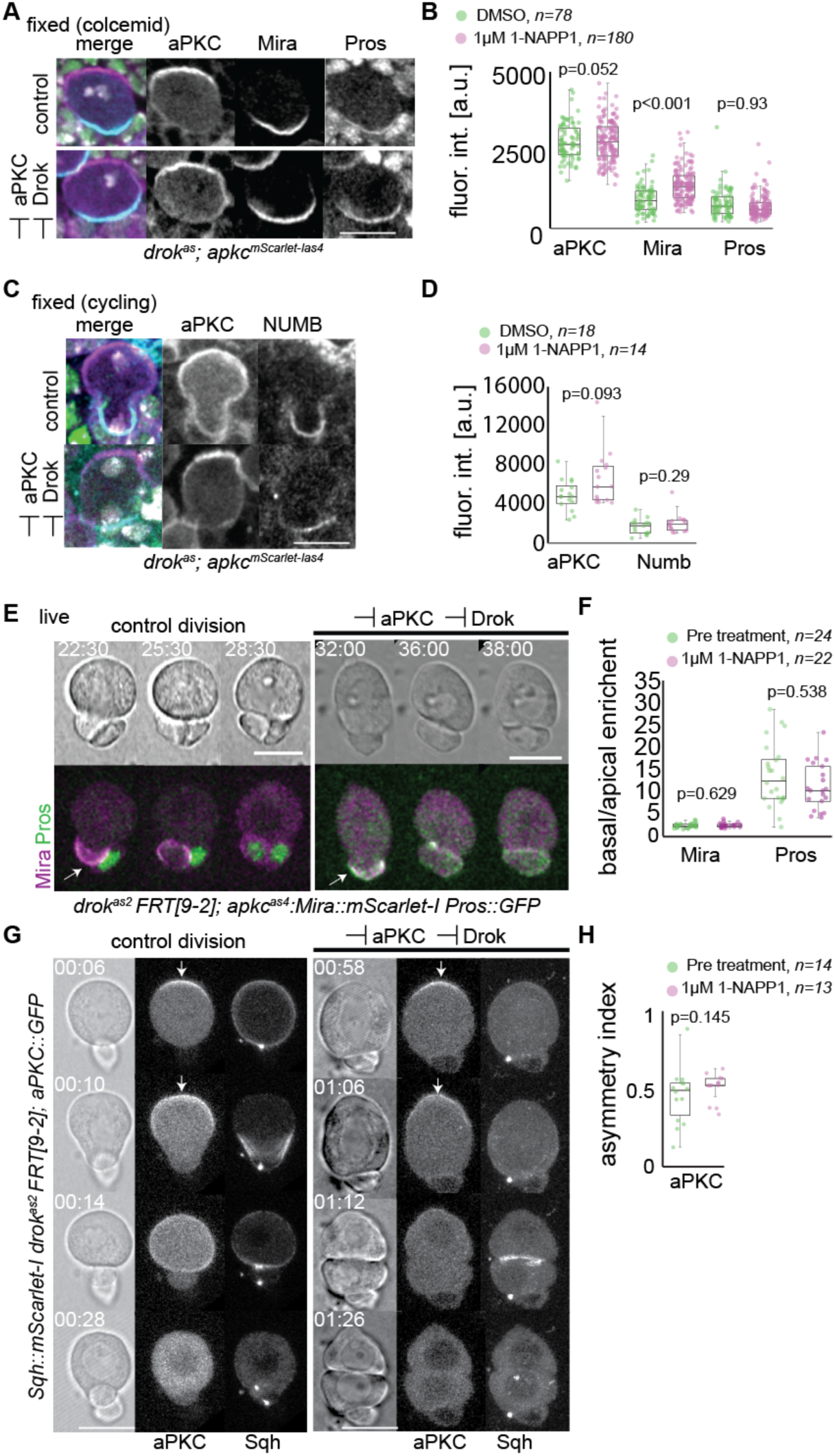
Fate determinants segregate asymmetrically upon partial co-inhibition of aPKC and Drok. **A**) *drok^as2^; apkc::mScarlet-I^as4^* brains were treated for 45min with 1µM 1-NAPP1 or DMSO (control) and 50µM Colcemid before being fixed and stained as indicated. **B**) Apical and basal fluorescence intensities at metaphase for aPKC as well as Mira and Pros. **C**) *drok^as2^; apkc::mScarlet-I^as4^* brains were treated for 45min with 1µM 1-NAPP1 or DMSO (control) fixed and stained as indicated. **D**) Apical and basal fluorescence intensities at telophase for aPKC and Numb, respectively. **E**) drok^as2^ FRT[9-2]/Y; apkc^as4^; Mira::mScarlet-I3 Pros::GFP larval neuroblasts in primary cell culture in the absence (control) or presence of 1µM 1-NAPP1. Mira and Pros segregate asymmetrically to the basal daughter in both cases (arrows). **F**) Basal and apical cortical intensities for Mira and Pros were background subtracted and normalised to background subtracted cytoplasmic values at anaphase. At the basal pole, Mira intensity was ∼ two-fold enriched over the apical pole values in the control and upon 1µM 1-NAPP1. Pros remained about 10-fold enriched in both cases. **G-H**) Two successive divisions of a larval neuroblast in primary cell culture in which aPKC segregates to the apical cell (arrows). **G**) Localisation of aPKC and Sqh in controls and upon addition of 1µM 1-NAPP1. **H)** Asymmetry index of aPKC for controls and partial Drok and aPKC co-inhibition. Scale bars: 10µm. Statistical tests: Mann-Whitney U test.

However, asymmetry of Mira as well as Pros was not significantly altered upon partial co-inhibition of aPKC and Drok in cultured larval neuroblasts (**Figure 5E,F, MOV7**). Live imaging further showed that aPKC remained asymmetrically localised during divisions with strongly reduced size asymmetry upon partial co-inhibition of aPKC and Drok (**Figure 5G,H**) whereas Sqh appeared reduced at both the cortex and the cytokinetic ring (**Figure 5G; MOV8**). Mira and Baz nevertheless segregated asymmetrically (in 26/26 of such divisions analysed Baz formed apical crescents) during such divisions, with each determinant inherited by only one daughter cell (**MOV9**).

Thus, partial co-inhibition of Drok and aPKC can severely alter GMC size while largely preserving neuroblast polarity and asymmetric fate-determinant segregation.

### Daughter-cell size predicts GMC versus neuroblast-like behaviour

Having established an experimental condition in which daughter-cell size asymmetry was strongly altered while fate-determinant segregation remained asymmetric, we next asked whether the size of the determinant-inheriting daughter affected its subsequent behaviour. To distinguish the consequences of altered daughter-cell size from continued kinase inhibition, we transiently co-inhibited Drok and aPKC in primary neuroblast cultures with 1µM 1-NAPP1 for 1h, washed out the inhibitor, and followed daughter-cell behaviour by long-term live imaging for up to 15h. Transient co-inhibition generated larger apical daughters inheriting the PAR complex (termed ‘cell 1’) and smaller basal, daughter (termed ‘cell 2’) which spanned a broad range of sizes, allowing us to relate their size to subsequent behaviour. Given that under partial inhibition of aPKC and Drok, fate determining molecules segregate asymmetrically (**Figure 5**), it follows that basal cells (cell 2) inherit fate determinants.

Strikingly, the size of cell 2 correlated with its behaviour after washout (**Figure 6A–D, MOV10**). Daughter cells with a measured cross-sectional area below 55µm² underwent no more than one subsequent division during the 15h observation period. When those cells divided, they appeared to do so in a size-wise symmetric manner and displayed long cell cycles of at least 6.6h and up to 11.1h. By contrast, daughter cells larger than 55µm² underwent at least three successive divisions, with some dividing up to seven times within the same observation period. These divisions displayed a size-asymmetry and GMC sizes resembling controls (**MOV10**), and their cell-cycle durations were shorter than 4.7h and more closely resembled those of control neuroblasts.

**Figure 6.**
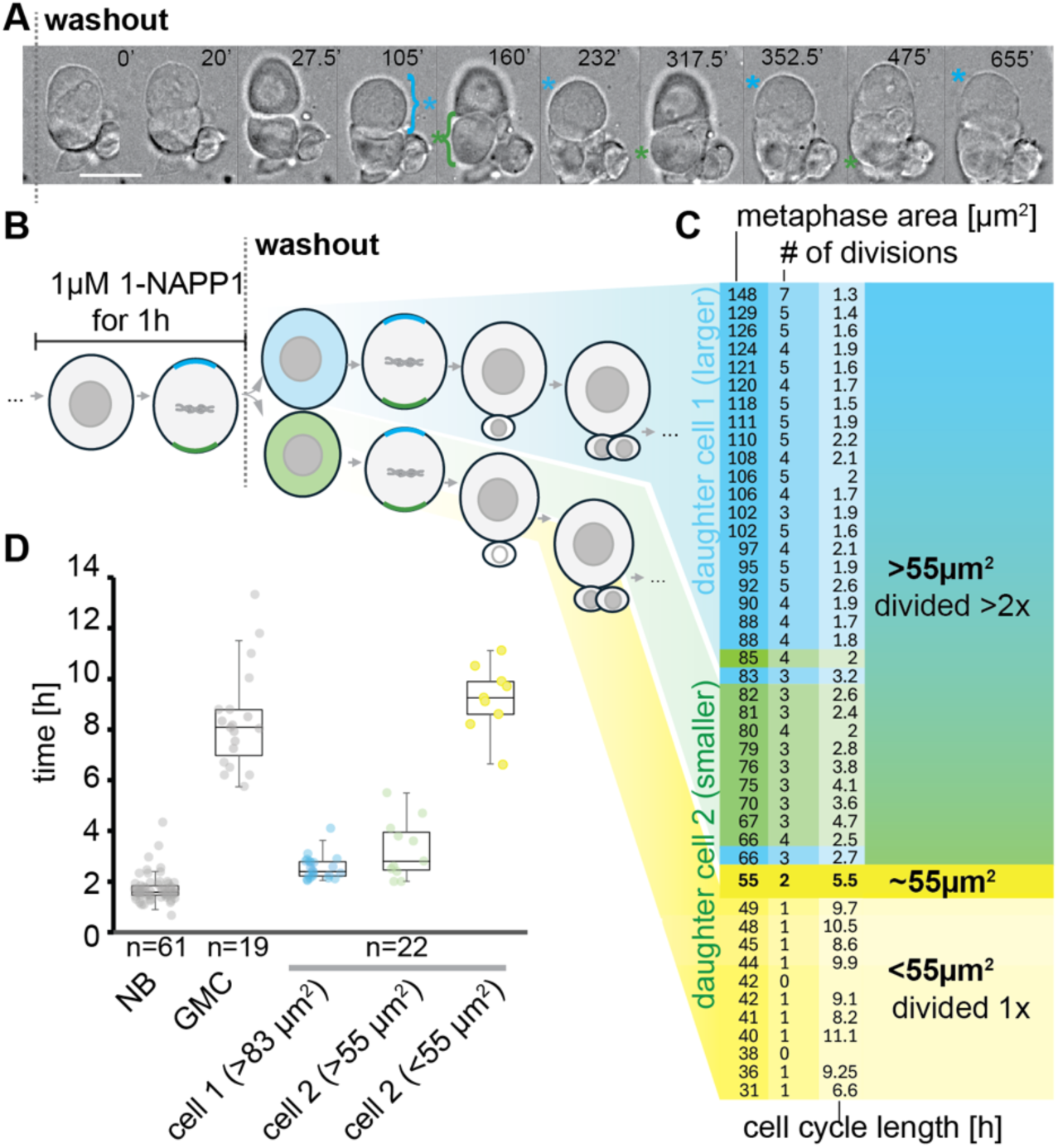
GMC-like behaviour is restricted to daughters below a cell-size threshold. **A**) A *drok^as2^ apkc^as4^* double mutant larval neuroblasts in primary cell culture treated for 1h with 1µM 1-NAPP1 divided ∼20min after inhibitor washout in a near symmetric manner (see corresponding **MOV10**). Both daughter cells resulting from this division continued to divide repeatedly (blue and green asterisks, respectively) in a size-wise asymmetric manner; scale bar:10µm, time in minutes. **B)** Schematic of experimental design and phenotypic outcomes. **C)** Upon acute and partial co-inhibition the cell 1 is defined as the larger apical daughter cell, cell 2 as the smaller basal daughter that inherited the fate determinants. Below a certain size cell 2 daughters underwent a maximum of one division, with long (>6.6h) cell cycles in a 15h experimental time window. Above this volume, they continued to divide asymmetrically at least three times with a maximum of seven times in the same 15h time window with cell cycle times <4.7h. When cell 2 did not divide, the last interphase area was measured. **D**) Quantification of cell cycles of control *w^1118^* neuroblasts treated with the same regime and of the larger cell 1 and the smaller cell 2 resulting from divisions with defective cell size asymmetry upon temporal, partial Drok and aPKC co-inhibition.

Thus, after transient co-inhibition, GMC-like proliferative behaviour was restricted to daughters below ∼55µm², whereas larger determinant-inheriting daughters retained neuroblast-like division dynamics. We therefore conclude that the small size of the GMC contributes to determining its fate.

## DISCUSSION

Here, we identify daughter-cell size as a determinant of lineage behaviour in *Drosophila* neuroblasts. Acute manipulation of aPKC and Drok allowed daughter-cell size to be altered while asymmetric polarity and fate-determinant segregation were largely preserved. Following restoration of kinase activity, determinant-inheriting daughters above a certain size displayed neuroblast-like proliferative behaviour, whereas smaller daughters remained GMC-like. These findings reveal a functional contribution of daughter cell size to the GMC fate decision.

A broader outcome of this study is the establishment of two complementary approaches for acute inhibition of *Drosophil*a Rho kinase (**Figure 1**). Pharmacological manipulation of Myosin activity has been limited because Blebbistatin is ineffective in fly cells (Straight et al. 2003), whereas Y-27632 has a narrow specificity window, inhibiting multiple kinases at concentrations only one order of magnitude above its IC_50_ (Davies et al. 2000). Importantly, Y-27632 also inhibits aPKC in *Drosophila* complicating interpretation, particularly in polarity-dependent processes where Drok and aPKC may both contribute, such as epithelial tissue mechanics. The analog-sensitive *drok^as2^* allele provides a chemical-genetic solution. Although the M164A mutation reduces DrokCAT activity *in vitro* to ∼20% (**Figure 1 supplement 1**), homozygous *drok^as2^* flies remain viable and fertile. This reduction may therefore reflect effects of the mutation on the truncated DrokCAT construct, or mutant full-length analog-sensitive Drok may retain sufficient *in vivo* activity to support development. BAY549 provides a complementary pharmacological approach. Our results show that both BAY549 and the analog-sensitive strategy potently inhibit Drok *in vitro*, with IC_50_ values in the low nanomolar range (**Figure 1**).

A central feature of our approach is that severe changes in daughter-cell size can be generated without loss of molecular asymmetry. Strong Drok inhibition markedly reduced cortical Myosin and delayed cytokinesis (**Figure 2**), but did not abolish neuroblast polarity or asymmetric determinant localisation (**Figure 3**). Similarly, under partial aPKC and Drok co-inhibition, severe changes in daughter-cell size occurred while aPKC, Mira, Pros and Numb remained asymmetric (**Figure 5**). Thus, this approach allows physical and molecular asymmetry to be uncoupled. It is currently unclear why basal Mira levels increase during Colcemid arrest (**Figure 5**); one possibility is that prolonged metaphase allows more efficient recruitment of Miranda from the cytoplasm. Importantly, unlike chronic genetic manipulations, the acute, transient and partial inhibition of aPKC and Drok used here restricts the perturbation to a short temporal window. This is expected to limit compensatory responses and cumulative pleiotropic or secondary effects associated with prolonged disruption of these kinases. Overall, our results therefore suggest that these manipulations largely preserve daughter-specific molecular segregation while altering the physical boundary between daughter cells.

We found that a daughter cell with a cross-sectional area of ∼55µm² displayed an intermediate phenotype, undergoing two divisions with a cell-cycle duration of ∼5.5h. This argues against an arbitrary separation of GMC- and neuroblast-like behaviours and instead suggests a transition in proliferative potential around this size. Smaller daughters (<49µm²) remained restricted to a more GMC-like programme with long cell cycles (>6.6h), whereas progressively larger daughters displayed increasingly neuroblast-like division dynamics, i.e. multiple divisions with shorter cell cycle length (**Figure 6**). Our data therefore suggest a transition in behaviour around 55µm² in this dataset that predicts subsequent lineage behaviour hinting at a size-dependent fate switch. Daughters below this threshold underwent at most one division and displayed long cell cycles, whereas larger daughters divided repeatedly, shortened their cell cycles and re-established asymmetric daughter-cell production (**Figure 6C,D**). Strikingly, these neuroblast-like properties most likely emerged despite inheritance of GMC fate determinants during the preceding division.

These findings indicate that inheritance of GMC determinants alone is not sufficient to impose GMC-like behaviour when daughter-cell size remains above an apparent threshold. Instead, cell size appears to modulate the ability of inherited fate determinants to enforce differentiation, providing a potential mechanism by which physical asymmetry during cytokinesis contributes directly to lineage specification. Delgado et al. (2025) similarly show that disrupting cell-size asymmetry increases the neuroblast pool, supporting an instructive role for size in lineage outcome (Delgado et al. 2025). Their chronic perturbations produce smaller neuroblasts with reduced lineage expansion and fewer differentiating progeny. Our acute, transient and reversible manipulation separates daughter-cell size from persistent perturbation of contractility or spindle geometry more accurately and identifies a threshold above which determinant-inheriting daughters retain neuroblast-like behaviour. Together, these studies suggest that cell size both influences stem-cell identity and constrains proliferative capacity during neuroblast lineage progression *in vivo.* It would be interesting in the future to ascertain the impact on neuroblast temporal identities.

An intriguing parallel may exist in type II neuroblast lineages, where the smaller daughter generated by the neuroblast becomes an intermediate neural progenitor (INP) rather than a terminally dividing GMC (Bowman et al. 2008; Bayraktar et al. 2010). Quantitative live imaging showed that INPs are approximately 1.5-fold larger than type I GMCs and, strikingly, mature INPs are also about 1.5-fold larger than the GMCs they themselves generate as type I and type II GMCs are similar in size. INPs additionally increase in size by approximately 1.7-fold during their maturation period (Homem et al. 2013). These observations raise the possibility that daughter-cell size contributes more generally to setting proliferative potential, with the larger size of INPs helping to permit limited self-renewal while their smaller GMC daughters become restricted to terminal division.

We identified aPKC as a regulator of GMC size (**Figure 4A,B**). aPKC may influence the cortical polarity framework within which Drok-dependent Myosin activity generates asymmetric cortical forces (Roubinet et al. 2017; Tsankova et al. 2017). Stronger inhibition of aPKC alone significantly enlarges GMCs, whereas partial inhibition of either aPKC or Drok individually does not (**Figure 4**), suggesting partially overlapping contributions to cortical mechanics and furrow positioning. Partial inhibition of either pathway may preserve sufficient spatial asymmetry and Myosin activity for near-normal daughter-cell size, whereas simultaneous inhibition could weaken both spatial organisation and mechanical execution of asymmetric cytokinesis. The altered recruitment and dissociation dynamics also suggest that aPKC depends at least in part on Drok activity (**Figure 3B’**).

Mechanistically, aPKC may modulate components of the established cell-size asymmetry machinery. Baz/aPKC and Pins/Gαi act in parallel to regulate spindle geometry and daughter-cell size (Cai et al. 2003). Spindle-derived CPC– centralspindlin/Tumbleweed cues and Sds22-dependent spindle positioning regulate Myosin redistribution (Roubinet et al. 2017), whereas Par3/Pins recruit PsGEF– RhoGAP54D to promote apical Myosin clearance (Loyer et al. 2026b) to which Pkn also contributes (Tsankova et al. 2017). aPKC could therefore phosphorylate these or other components linking cortical and spindle-derived pathways for cell size asymmetry, coordinating their activity with the polarity axis. Identifying such substrates will clarify how aPKC contributes to physical asymmetry and links cell-size asymmetry to cell polarity in neuroblasts. aPKC has been proposed to promote neuroblast self-renewal, as reducing aPKC decreases neuroblast numbers (Rolls et al. 2003; Lee et al. 2006). Our findings suggest a potential mechanistic explanation for this: enlarged GMC production may progressively reduce neuroblast size, ultimately compromising self-renewal eventually causing a reduction of neuroblast number in line with the conclusions from Delgado et al.

Cell size could influence fate by changing the effective concentration and stoichiometry of inherited determinants in line with previous conclusions from studies regarding the role of spindle orientation in neuroblasts (Cabernard and Doe 2009). If similar amounts of Pros, Brat and Numb enter daughters of different volumes, enlargement could reduce Pros nuclear concentration, weaken Brat relative to its RNA targets, and lower Numb activity relative to the Notch machinery, collectively suppressing the GMC differentiation programme. Alternatively, GMC fate might depend on the ability to inherit as little cytoplasm from the neuroblast as possible. In this scenario the neuroblast cytoplasm might contain yet to be identified stemness or self-renewal factors. This would be in line with the observation that neuroblasts retain polarity and repeated asymmetric divisions after isolation from their native tissue environment (Homem et al. 2013), supporting a substantial cell-intrinsic component to neuroblast fate.

Together, our study has generated novel tools for specific Rho kinase inhibition in *Drosophila*. Our findings further suggest that eccentric cytokinesis does more than partition fate determinants: by controlling the amount of cytoplasm inherited by each daughter, it also establishes a physical context in which those determinants can act. Cell size may therefore represent an intrinsic component of cell-fate specification in this context, linking division geometry to the proliferative potential of daughter cells.

## MATERIALS AND METHODS

### Fly stocks and genetics

Flies were reared on standard corn meal food at 25 °C.

**Figure 1 genotypes**

*sqh::GFP; apkc^as4^* (*apkc^as4^*)

sqh::mScarlet-I ^BL#94929^ *drok^as2^FRT[9-2]*

*drok^as2^; apkc::mScarlet-I^as4^*

*drok^as2^*

**Figure 1 supplement 2 genotypes**

*drok^as2^; Ubi-PH::GFP; Ubi-His::RFP*

**Figure 2 genotypes**

sqh::mScarlet-I ^BL#94929^ *drok^as2^FRT[9-2]*

**Figure 2 supplement genotypes**

Histone2A::RFP; CD4:GFP

**Figure 3 genotypes**

sqh::mScarlet-I ^BL#94929^ *drok^as2^FRT[9-2]; aPKC::GFP*

*drok^as2^; Mira::mCherryHA*

*drok^as2^; Numb::GFP; Ubi-His-RFP*

**Figure 4 genotypes**

*apkc::mScarlet-I^as4^*

**Figure 5 genotypes**

*drok^as2^FRT[9-2]; apkc::mScarlet-I^as4^*

*drok^as2^ FRT[9-2]/Y; apkc^as4^; Mira::mScarlet-I3 Pros::GFP*

*sqh::GFP; apkc^as4^ apkc::mScarlet-I^as4^*

**Figure 6 genotypes**

*drok^as2^FRT[9-2]; apkc::mScarlet-I^as4^*

### Movie genotypes

**Movie 1**. *w^1118^* (BL#53502), apkc^as4^ (Hannaford et al. 2019)

**Movie 2.** drok^as2^/Y; worniu-Gal4 (Albertson et al. 2004) UAS-nlsMCP::GFP (Jayanandanan et al. 2011) UAS-mCherry::Jupiter (Chabu and Doe 2008)

**Movie 3.** sqh::mScarlet-I ^BL#94929^ *drok^as2^FRT[9-2]*

**Movie 4**. *drok^as2^FRT[9-2]; numb::GFP (CrispR knock in, gift from Roland Le Borgne)*

**Movie 5.** drok^as2^FRT[9-2]; apkc^mScarlet-Ias4^

**Movie 7.** drok^as2^ FRT[9-2]/Y; apkc^as4^; Mira::mScarlet-I3 Pros::GFP

**Movie 6.** drok^as2^FRT[9-2]; apkc^mScarlet-Ias4^

**Movie 8.** Sqh::GFP (Ambrosini et al. 2019); *apkc^mScarlet-I as4^*

**Movie 9**. Baz::GFP (BL#51572); *apkc^as4^*, Mira::mCherry (Ramat et al. 2017)

**Movie 10** drok^as2^FRT[9-2]; apkc^mScarlet-Ias4^

### Sources of fly lines

*w^1118^* (BL#53502)

*sqh::GFP* (Suzanne lab, University of Toulouse, (Ambrosini et al. 2019))

*apkc^as4^* (Hannaford et al. 2019) sqh::mScarlet-I (Parkhurst lab, BL#94929) *drok^as2^* (this study)

*drok^as2^FRT[9-2]* (this study, recombinant made by Luschnig lab, University of Münster)

*aPKC::GFP* (StJohnston lab, University of Cambridge(Erdmann et al. 2019))

*Numb::GFP* (LeBorgne lab, University of Rennes (Bellec et al. 2018))

Ubi-His-RFP (Schuh et al. 2007)

*Pros::GFP* (Song lab, Peking University (Liu et al. 2020))

*Mira::mScarlet-I3* (this study)

Mira::mCherry (Ramat et al. 2017) Baz::GFP (BL#51572)

UAS-mCherry::Jupiter (Chabu and Doe 2008)

worniu-Gal4 (Albertson et al. 2004)

Ubi-PH::GFP (Gervais et al. 2008) Histone2A::RFP

CD4:GFP

### Generating the analog sensitive drok^as2^ allele

To generate an analog sensitive DROK version the gatekeeper residue Methionine 164 in CG9774-PA (FBpp0074061) was changed to Alanine by genome editing using CrispR resulting in the *drok^as2^* allele. This guide RNA (taaagtccatcaccatgtatagg) and oligodeoxynucleotide(atgtcgtagtcacccatcagcgagactatatcgccgccgggcataaagtcCGCcaccatgtata ggtatttggcatcctgtgagaaaagattttatgaattaatt) were injected into yw;;nos-Cas9(III-attP2, BL78782) and successful editing was confirmed by PCR on genomic DNA using these primers (5’ aaatatcattgttagcccact and 5’ attcgcatacaggtgccaa) followed by sequencing. *drok^as2^* flies are homozygous viable and fertile, since *drok^as2^* was generated by CrispR it was recombineered to FRT[9-2] to label the chromosome with a traceable marker (w+) to more easily facilitate genetic crosses and further recombination.

### Generating mira^mScarletI-3^

Mira::mScarletI-3 was generated via CrispR. A 22 amino acid linker, (5x(GGGS) and an XbaI site followed by the mScarletI-3 sequence and a R S I T S Y N V C Y T K L S A S peptide linker sequence was inserted after Q700 of Mira PA. The resulting *mira^mScarletI-3^* flies are homozygous viable and fertile.

### Immunostainings

Larval brains were dissected in phosphate buffered saline (PBS) and fixed in 4% Formaldehyde (#F8775, Sigma) for 20 min at room temperature (RT), tissues were permeabilised in for 2 hr in PBS-Triton 0.1% (PBT) at RT. They were then rinsed twice in PBT, washed for 10 minutes in PBT, rinsed twice again in PBS, placed in 50% glycerol (#49781, Sigma) and finally mounted in Vectashield (#H-1000, Vector Laboratories).

### Antibodies

#### Primary antibodies

Phospho-MYL9 raised in mouse (Ser19, Invitrogen, P-Sqh)

Anti-Numb raised in sheep (Loyer et al. 2024)

anti-Mira raised in sheep (Loyer et al. 2024)

Anti-Pros raised in mouse (MR1A, DSHB)

#### Secondary antibodies

Donkey anti-Sheep IgG Alexa 546 (#A-21098, Thermofisher)

Donkey anti-Sheep IgG Alexa 488(#A-11015, Invitrogen)

Donkey anti-Mouse IgG Alexa 488 (#A32766, Thermlife Tech)

DAPI solution (1 mg/mL #MBD0015-1ML, SIGMA)

Alexa Fluor™ 647 Phalloidin (#A22287, Invitrogen)

### Neuroblast primary culture and drug treatment

Schneider’s medium (SLS-04–351Q) supplemented with glucose (1 mg/ml), FCS 10% (#A5670501, ThermoFisher), fly extract 2.5% (DGRC) and insulin 0.07 mg/ml (#12585014, ThermoFisher) was used to culture larval neuroblasts. L3 brains were dissected in collagenase buffer (800mg NaCl, 20mg KCl, 5mg NaH2PO4, 100mg NaHCO3, 100mg glucose in 100ml distilled water) incubated in that buffer containing 0.2 mg/mL collagenase (#C0130-100MG, Sigma) for 15 minutes, rinsed in collagenase buffer and transferred into Schneider’s medium with 10mg/ml Fibrinogen (Januschke and Loyer 2020). A drop of 8 µl of Schneider’s medium with Fibrinogen containing the brains was pipetted onto a Poly-L-Lysin-coated glass dish (#FD35PDL-100, World Precision Instruments). Brains were manually dissociated within the drop. The cells were allowed to settle for ∼5 minutes, after which clotting was induced by adding 1 µl of Thrombin 100 units/ml (#T4648-1KU, Sigma) on 4 sides of the Schneider and Fibrinogen drop. Clots were covered with 200µl of supplemented Schneider’s medium. 1-Naphthyl PP1 (1-NAPP1, #T3935-10mg, Chempur.de) or BAY549 (#TC-S 7001, Tocris) was added at varying concentrations to the medium typically by adding twice the desired final concentration in 200µl supplemented Schneider’s medium to the 100l already present. Washout was performed by repeatedly pipetting off 350µl of medium and replacing it for at least 10x without perturbing the clot.

### Live imaging, fluorescent signal measurement and image processing

Primary neuroblasts in culture were either imaged on an inverted Olympus IX81 microscope equipped with a 60x, 1.42NA PlanApoN objective, a Coolsnap HQ2 camera (Photometrics), an automated XY stage with a piezo top plate (Applied Scientific Instrumentation) and a DC4104 controller driving a 625nm LED (Thorlabs) all controlled by Micro-Manger (Edelstein et al. 2014). Or a LEICA SP8 Stellaris confocal microscope equipped with an 86x water immersion objective (NA 1.20) was used. All images were processed and analysed using ImageJ (Schneider et al. 2012). Rolling ball background subtraction with a radius of 50 pixel or 100 pixel and a gaussian blur with a sigma of 0.8 pixels were applied for figure panels, intensity measurements were carried out on raw images. We corrected the drifting of movies with our custom AutoHyperstackReg macro (Januschke and Loyer 2020), based on the TurboReg and MultiStackReg plugins (Thévenaz et al. 1998). Cytoplasmic intensity was measured inside manually drawn polygonal shapes. Measurement of cortical signal intensity was performed using our custom rotating linescans macro (Januschke and Loyer 2020). Background fluorescence was measured outside of neuroblasts and subtracted from any other measured signal. The asymmetry index (AI) was calculated using this formula: AI^apical^ =(value^apical^ – value^basal^)/(value^apical^ + value^basal^) or AI^basal^=(value^basal^ – value^apical^)/(value^apical^ + value^basal^).

### *In vitro* kinase assay

The kinase assay was essentially performed as described (Hannaford et al. 2019). Since the catalytically active fragment of vertebrate Rho kinase has been successfully used in *in vitro* kinase assays previously (Amano et al. 1996) we cloned the DNA sequence coding for amino acids 2-530 of Drok-PA (FBpp0074061, DrokCAT) into thepCMV5-Flag1 vector by Gibson assembly yielding pCMV5 FLAG DrokCAT^WT^. Site directed mutagenesis was used to generate the analog sensitive version pCMV5 FLAG DrokCAT^as2^ and the kinase dead version in which the DFG motif has been mutated to DAG yielding pCMV5 FLAG DrokCAT^KD^.

The apparent IC_50_ was determined under activity-matched conditions (40µg DrokCAT^WT^had about the same activity as 160µg DrokCAT^as2^). Peptide used: long S6 peptide (KEAKEKRQEQIAKRRRLSSLRASTSKSGGSQK). For the positive control His-ROCKII (DU19085) was used with a stock concentration of 0.49 mg/ml. All assays carried out for 30min at 30°C. The empty FLAG vector lysate was used as the background control for all the assay calculations. The empty FLAG vector control counts were slightly higher than the substrate peptide only control and this small window appeared inhibitable possibly explaining why some negative values were obtained for BAY549.

Four 15 cm plates with 12 × 10⁶ HEK293 cells per plasmid were transfected with 26µg plasmid and 78µl Polyethylenimine per plate. Cells were harvested 30.5h after transfection yielding ∼25mg of protein after lysis in lysis buffer (270mM sucrose 50mM Tris (pH 7.5) 1% Triton X-100, 1mM EGTA (pH 8.0), 1x PIC, 50mM NaF, 10mM β-glycerophosphate, 1 mM sodium orthovanadate, 5 mM sodium pyrophosphate). For lysis cells were scraped in 10 ml PBS/plate — and plates pooled per plasmid. Cells were centrifuged at 500g for 5in and the pellets lysed in 2ml lysis buffer and snap frozen. Assays using 1-NAPP1, BAY549 or Y-27632 (#1254/1, Tocris) were performed in triplicate, using 100uM ATP and incubated in kinase assay for 30 mins (after checking this was still in linear range). The empty FLAG vector lysate was used as the background control for all the assay calculations.

### Wing disc contraction assay

L3 larvae were dissected in Schneider’s media (#21720024, ThermoFisher). Multiple imaginal wing discs were mounted on 35mm glass bottom dishes (#FD35100, World Precision Instruments), using fibrinogen and thrombin as described for neuroblast cultures to fix the discs in place on the dish. According to the experimental condition, an aliquot of the dissection media was added to the discs, containing either the appropriate concentration of a given inhibitor, or an equivalent volume of the appropriate DMSO concentration. Compounds used in this assay: BAY-549, BAY-4900 (obtained from https://www.bayer.com/en/pharma/chemical-probes-open-access), Latrunculin B (Lat B, # L5288-1MG, Sigma), or DMSO (#D2653-5X5ML, Merck). Imaging was carried out using an Olympus IX81 microscope set up described above, with a 10x objective lens (UPlanSApo, 0.4NA). Disc were imaged every 60 seconds at a single z-slice with both bright field and fluorescent images collected at each time point. Imaging was carried out for a preliminary 10-minute period as a control before any further treatment. Following this, an additional aliquot of media was added, containing either another inhibitor, or 1NA-PP1 or DMSO of equivalent volume. A second round of imaging was then started, lasting 60 minutes. Image stacks were imported into FIJI software. Using the fluorescent images, a binary mask was produced for each time point, reflecting wing disc area change. Measurements were taken for each time point in the 60-minute imaging cycle. For MOVIE 1 wing discs were placed at the bottom of different wells of an IBIDI µ-Slide 8 Well (Cat.No: 80826) without fibrinogen to illustrate the contractions better.

### Quantitative data representation and statistical analysis

Design of boxplots: dots: individual measurements; centre line, median; box limits, upper and lower quartiles; whiskers, 1.5×interquartile range. p Values were calculated using a non-parametric two-tailed Mann–Whitney U test unless indicated otherwise.

## Supporting information

MOV1

MOV2

MOV3

MOV4

MOV5

MOV6

MOV7

MOV8

MOV9

MOV10

## Acknowledgements

We thank M. Suzanne, D. St Johnston, R. LeBorgne, F. Schweisguth, Y. Song. C. Doe and M. Leptin for sharing flies. We thank the Dundee imaging facility for excellent support, L. Plater and R. Tooth for excellent technical support and S. Luschnig for recombining *drok^as2^* onto FRT[9-2]. We thank C. Gonzalez and C. Cabernard for discussion. Stocks obtained from the Bloomington Drosophila Stock Centre (NIH P40OD018537) were used in this study. This work was supported by grants from the BBSRC (BB/V001353/1 and BB/T017546/1) and by the Wellcome Trust grant 218520/Z/19/Z supporting a PhD studentship to IJM. For the purpose of open access, the authors have applied a Creative Commons Attribution (CC BY) licence to any Author Accepted Manuscript version arising from this submission.

## Author contribution

JJ and IJB conceived the study. JJ designed most experiments. IJB, NL, HM, CR designed experiments and IJB, NL, MD, IM, HM and CR carried out experiments. IJB, NL, CR and JJ analysed and interpreted the data. JJ and KD acquired funding, JJ supervised the project and wrote the manuscript that was agreed upon by all authors.

*The authors declare no competing interest*.

## SUPPLEMENTARY FIGURES

**Figure 1 supplement 1.**
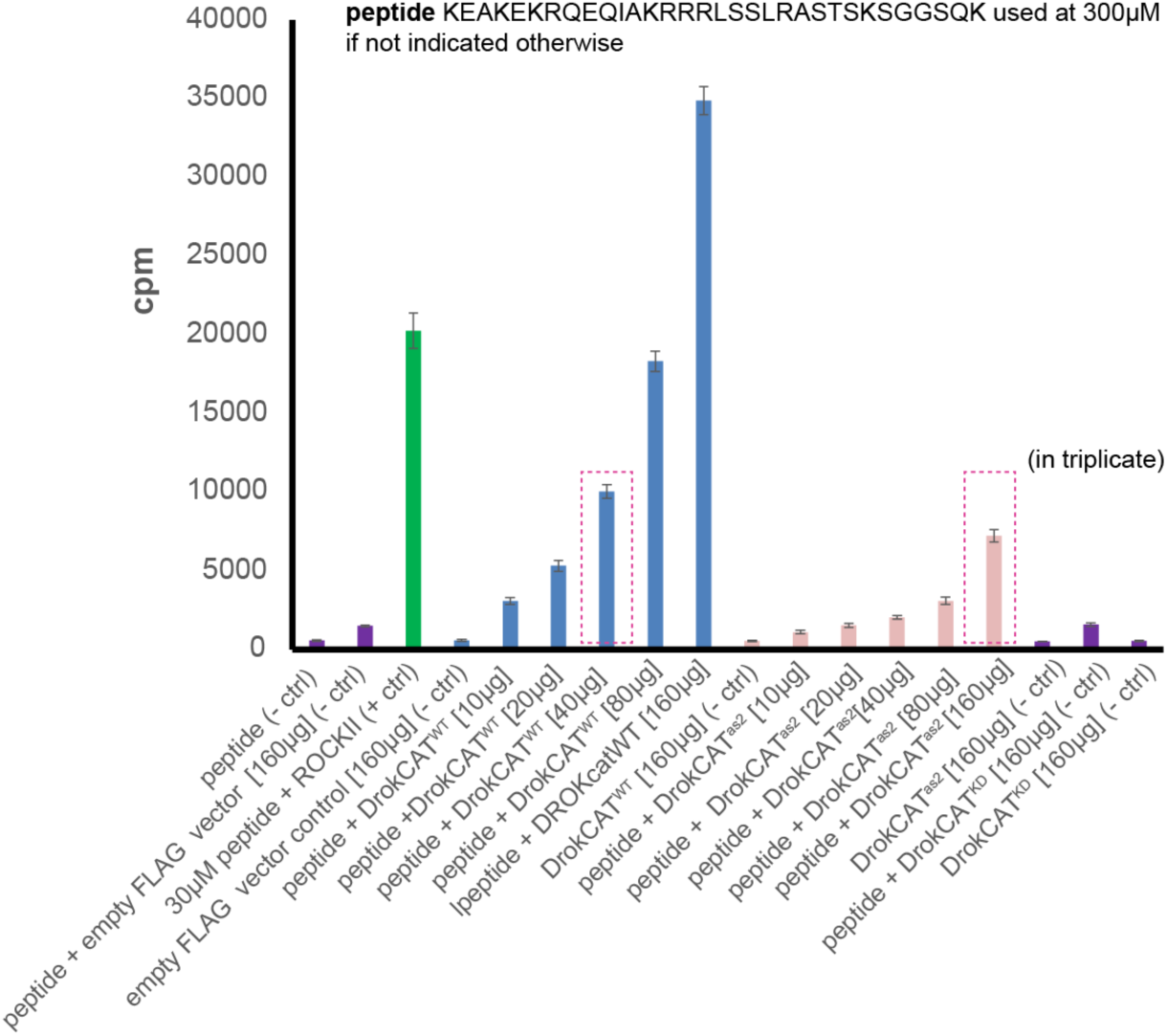
Basis for activity matching of DrokCAT^WT^ and DrokCAT^as2^. The constructs (see methods) were transfected into HEK293 cells and immunoprecipitated from lysates using the FLAG-tag. Increasing amounts of the lysates were tested for activity. Of note DrokCAT^as2^ has approximately 20% of DrokCAT^WT^ activity without the inhibitor added. The peptide used in the assay is indicated above, we estimated that 160µg of *DrokCAT^as2^* lysate has the best comparable the activity to 40µg DrokCAT^WT^ lysates (dashed magenta boxes). The kinase dead (KD) construct and all other negative controls have neglectable activity. Note that 30µM of the peptide were used in the positive control using ROCKII kinase. [µg] indicates the amount of lysate used.

**Figure 1 supplement 2.**
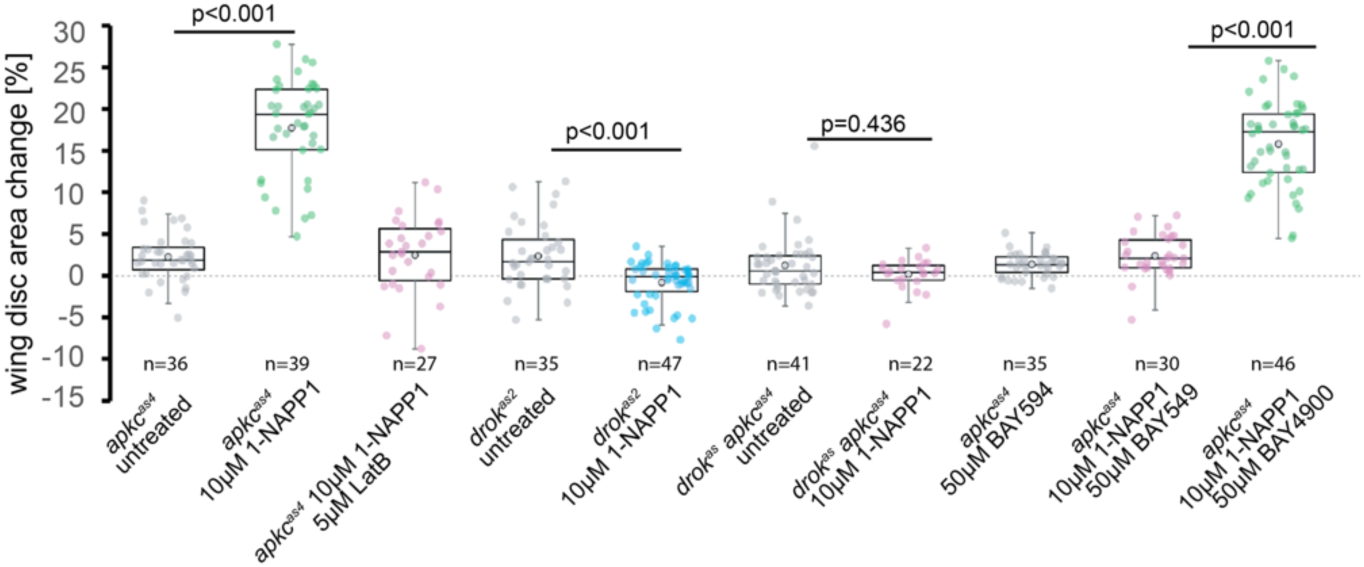
drok^as2^/1-NAPP1 and BAY549 disrupt Myosin-dependent contractility in vivo. Wing disc contractility suppression assay measuring the ability of the indicated means to inhibit Drok to suppress contractions of wing discs in culture induced by aPKC inhibition. Apical constriction of epithelial cells depends on Rho kinase activity (Sawyer et al. 2010), and acute inhibition of aPKC using the *apkc^as4^* allele induces ectopic apical constriction and contraction of eye imaginal discs (Hannaford et al. 2019). We reproduced this phenotype in wing discs (**MOV1**) and used it to confirm Drok inhibition *in vivo.* Depolymerisation of actin with Latrunculin B (Lat B) suppressed such disc contraction, confirming that the response to aPKC inhibition was actin dependent. Co-inhibition of aPKC and Drok, using either BAY549 and 1-NAPP1 on *apkc^as4^* mutants or 1-NAPP1 on *drok^as2^*; *apkc^as4^* double mutant tissues similarly suppressed ectopic wing-disc contraction. In addition, inhibition of Drok alone with *drok^as2^*/1-NAPP1 caused a significant increase in disc area, consistent with tissue relaxation following reduced Rho kinase activity (Uehata et al. 1997).

**Figure 2 supplement 1.**
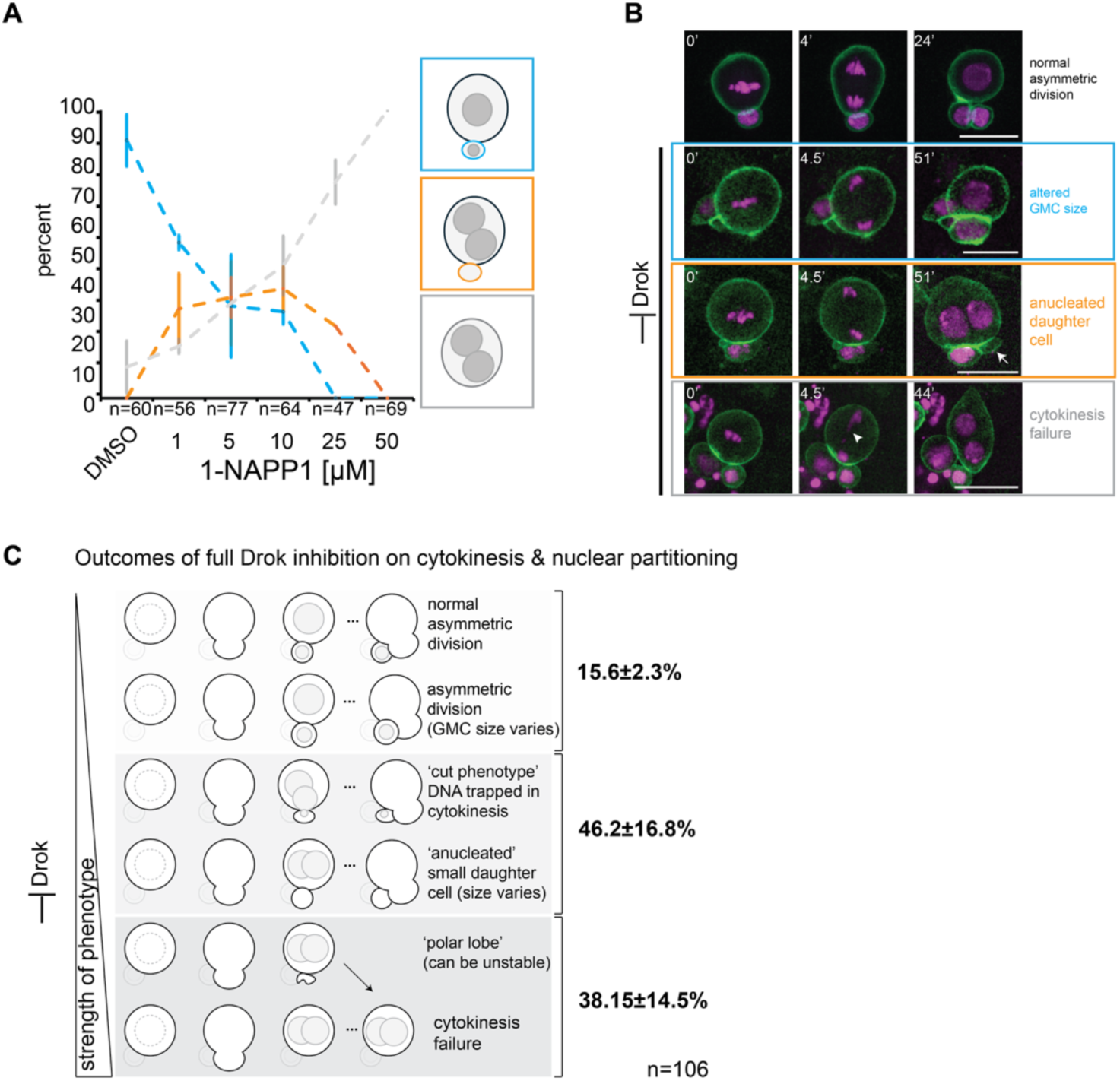
Strong Drok inhibition leads to cytokinesis and nuclear partitioning defects. **A**) Response of neuroblasts in primary cell culture observed by wide field microscopy to increasing concentrations of 1-NAPP1 resulting in dose dependent differences in the frequency of size wise asymmetric neuroblasts divisions, cytokinesis failure or binucleated neuroblasts with a productive cytokinesis generating an anucleated daughter cell. **B**) *drok^as2^*; PH::GFP; His::RFP larval neuroblasts in primary cell culture. Top panel, untreated (control), dividing normally. Below different outcomes of the effect of 10µM 1-NAPP1 on cytokinesis and nuclear partitioning. Such divisions can generate stable cells lacking a nucleus (arrow). Lagging chromosomes can be observed in neuroblasts that subsequently fail cytokinesis (arrowhead). Scale bar: 10µm. Time in minutes. **C**) Schematic summary of the observed phenotypes and the frequency at which they occurred. We observed cases in which the DNA of the nucleus destined for the basal cell got trapped or cut (termed ‘cut phenotype’) by the cleavage furrow resulting in what appeared successful cytokinesis. We also observed smaller patches of basal membrane that resembled polar lobes (Cabernard et al. 2010) that sometimes were absorbed or formed stable structures in other cases.

**Figure 2 supplement 2.**
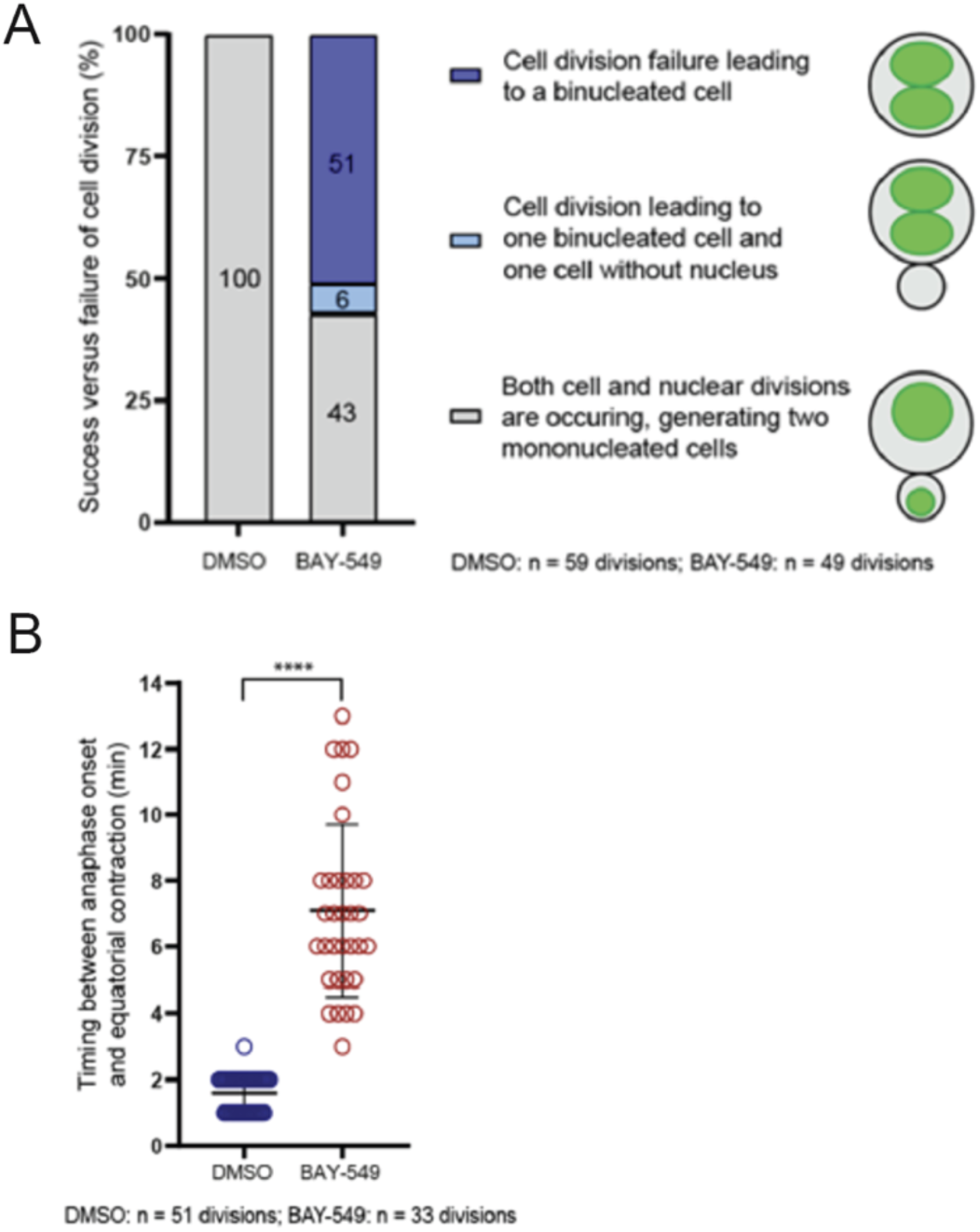
Effect of Drok inhibition by BAY549 on neuroblasts in whole mount brains. **A**) Percentage of neuroblasts in whole mount Histone2A::RFP and CD4:GFP expressing neuroblasts succeeding in cell division, failing in cytokinesis, or generating one mononucleated cell together with one anucleate cell. Larval brains were treated with either DMSO as control (n=59 mitotic cells) or 10µM of BAY-549 (n=49 mitotic cells). Data are compiled from at least three independent experiments per condition. **B**) Timing of equatorial contraction relative to anaphase onset, in minutes, in DMSO- or BAY-549-treated brains (n=51 and 33 mitotic cells, respectively). Bars indicate mean ± standard deviation. Asterisks denote statistical significance, derived from unpaired t tests: ∗∗∗∗ p ≤ 0.0001. Data are compiled from at least three independent experiments per condition.

## VIDEO LEGENDS

**MOVIE 1.** *Wing disc contractility assay*. Left *w^1118^* L3 larval wing disc, right apkc^as4^ wing disc both treated with 10µM 1-NAPP1. Time stamp: hh:mm.

**MOVIE 2**. *Generation of an anucleated daughter cell with a cortical microtubule network upon strong Drok inhibition*. A neuroblast in primary culture from a male *drok^as2^*; MCPnlsGFP mCherry::Jupiter larvae. Scale bar: 10µm. Time stamp hh:mm.

**MOVIE 3**. *Drok inhibition reduces cortical Myosin accumulation and slows the contraction of the cytokinetic ring*. A *Sqh::GFP drok^as2^ FRT[9.2]* neuroblast in primary cell culture prior and upon addition of 10µM 1-NAPP1. The outcome of this division is cytokinetic failure. Scale bar: 10µm. Time stamp hh:mm.

**MOVIE 4***. Numb segregates into the anucleated cell produced upon Drok inhibition*. A drok^as2^ FRT[9-2]; numb::GFP; Histone::RFP neuroblast in primary cell culture. Numb polarises in division and segregates to the smaller anucleated daughter cell. Scale bar: 10µm. Time stamp hh:mm.

**MOVIE 5.** *Partial co-inhibition of Drok and aPKC yields basal daughter cells of different sizes*. *drok^as2^FRT[9-2]; apkc^mScarlet-Ias4^* neuroblasts in primary cell culture. Left panel, DMSO treated control. The other three panels were treated with 1µM 1-NAPP1. Apical cell (cell 1) (all apical poles oriented to the top) outline with green dots, basal cell (cell 2) outlined with magenta dots. Colour bars indicate the apical to basal ratio at the end of the movie taken from the long axis of the cells. Time stamp hh:mm.

**MOVIE 6.** Near symmetric divisions of neuroblasts upon partial co-inhibition of Drok and aPKC. Three *drok^as2^FRT[9-2]; apkc^mScarlet-Ias4^* neuroblasts in a primary cell culture incubated with 1µM 1-NAPP1. Time stamp hh:mm.

**MOVIE 7**. *Mira and Pros segregate asymmetrically in controls and upon partial co-inhibition of Drok and aPKC.* A *drok^as2^ FRT[9-2]/Y; apkc^as4^; Mira::mScarlet-I3 Pros::GFP* neuroblasts untreated (top panels) or treated with 1µM 1-NAPP1 (Bottom panels). Scale bars: 10µm. Time stamp mm:ss

**MOVIE 8**. *aPKC polarises normally during a near symmetric division upon partial co-inhibition of Drok and aPKC.* A *Sqh::GFP; apkc^mScarlet-I as4^* neuroblast treated with 1µM BAY549 and 1µM 1-NAPP1. Left panel transmitted light to show cell outline. Right panel aPKC (magenta) and Sqh (green). Time stamp hh:mm.

**MOVIE 9.** Baz and Mira segregate asymmetrically during a near symmetric division upon partial co-inhibition of Drok and aPKC. A Baz::GFP; *apkcas4*; Mira::mCherry neuroblast treated with 1µM BAY549 and 1µM 1-NAPP1. Time stamp hh:mm.

**MOVIE 10.** Both daughter cells continue asymmetric neuroblast-like divisions after an initial near symmetric division induced by partial co-inhibition of Drok and aPKC. A *drok^as2^FRT[9-2]; apkc^mScarlet-I as4^* neuroblast treated with 1µM 1-NAPP1. The focus shifts at each division to visualise the small daughter cells that are transiently outlined and labelled in green for the apical cell (cell 1) and in magenta for the basal cell (cell 2). Time stamp: hh:mm:ss.

